# RANKL signaling sustains primary tumor growth in genetically engineered mouse models of lung adenocarcinoma

**DOI:** 10.1101/142620

**Authors:** Julien Faget, Caroline Contat, Nadine Zangger, Solange Peters, Etienne Meylan

**Affiliations:** Swiss Institute for Experimental Cancer Research, School of Life Sciences, Ecole Polytechnique Fédérale de Lausanne, CH-1015 Lausanne, Switzerland.; Bioinformatics Core Facility, Swiss Institute of Bioinformatics, CH-1015 Lausanne, Switzerland.; Department of Oncology, Centre Hospitalier Universitaire Vaudois, University of Lausanne, CH-1011 Lausanne, Switzerland.

**Author notes:** **Corresponding author:** Etienne Meylan, Swiss Institute for Experimental Cancer Research, School of Life Sciences, Ecole Polytechnique Fédérale de Lausanne, Station 19, CH-1015 Lausanne, Switzerland.

**Keywords:** lung adenocarcinoma, RANK ligand, genetically engineered mouse models, preclinical trial

## Abstract

**Hypothesis:** Non-small cell lung cancer (NSCLC) is the leading cause of cancer mortality. Recent retrospective clinical analyses suggest that blocking the receptor activator of NF-κB (RANK) signaling pathway inhibits the growth of NSCLC and might represent a new treatment strategy.

**Methods:** *RANK* and *RANKL* expression in human lung adenocarcinoma was interrogated from publicly available gene expression datasets. Several genetically engineered mouse models were used to evaluate treatment efficacy of RANK-Fc to block RANKL, with primary tumor growth measured longitudinally using micro-computed tomography. A combination of RANKL blockade with cisplatin was tested to mirror an ongoing clinical trial.

**Results:** In human lung adenocarcinoma datasets, *RANKL* expression was associated with decreased survival and *KRAS* mutation, with the highest levels in tumors with co-occurring *KRAS* and *LKB1* mutations. In *Kras^LSL-G12D/WT^, Kras^LSL-G12D/WT^*; *Lkb1^Flox/Flox^* and *Kras^LSL-G12D/WT^*; *p53^Flox/Flox^* mouse models of lung adenocarcinoma, we monitored an impaired progression of tumors upon RANKL blockade. Despite elevated expression of RANKL and RANK in immune cells, treatment response was not associated with major changes in the tumor immune microenvironment. Combined RANK-Fc with cisplatin revealed increased efficacy compared to single agents.

**Conclusions:** RANKL blocking agents impair the growth of primary lung tumors in several mouse models of lung adenocarcinoma, and suggest that patients with *KRAS* mutant lung tumors will benefit from such treatments.

## Introduction

With over 1.5 million deaths yearly, lung cancer has become the leading cause of cancer-related mortality worldwide, with non-small cell lung cancer (NSCLC) representing 80-85% of cases^1^. Although new therapies are emerging to combat lung adenocarcinoma, the main histological subtype of NSCLC, their use is currently limited to a subset of molecularly selected patients, and no specific targeted therapy has demonstrated activity against tumors carrying the most frequently mutated proto-oncogene, *KRAS*^2^. One interesting approach for cancer treatment is to identify actionable membrane-bound or secreted ligand-receptor systems that act in the tumor environment to promote disease progression, which can be very selectively blocked through the use of small compounds or antibodies. One such example is the tumor necrosis factor (TNF) family member, receptor activator of NF-κB (RANK) ligand (RANKL) that signals through its receptor RANK, first discovered for communication between T cells and dendritic cells^3^. Through its receptor RANK, RANKL activates osteoclasts for bone resorption, promotes lymphocyte maturation and function, and enables mammary gland and secondary lymph node organogenesis^4–6^. A link between RANKL signaling and cancer has been established in recent years: first, and perhaps owing to its bone remodeling capabilities, the RANKL-RANK pathway facilitates bone metastasis formation^7, 8^; second, it promotes seeding of breast tumor cells into the lungs in a T-regulatory (Treg) cell-dependent manner^9^; third, it participates in progestin-dependent mammary tumor development^10, 11^. The importance of this signaling pathway in primary breast cancer is well documented, with *RANKL* mRNA being stabilized in response to progesterone, leading to increased RANKL protein expression that promotes tumor cell proliferation^12^ and epithelial-mesenchymal transition to foster tumor cell migration^13^. In contrast, much less is known about its contribution in primary tumors from other carcinomas such as lung cancer. Recently, in a large international clinical trial of patients with advanced solid tumors (excluding breast and prostate cancer) or multiple myeloma, denosumab, a fully human monoclonal antibody that binds RANKL and blocks RANKL-RANK interaction, demonstrated activity in preventing bone-metastasis related adverse events, such as fracture or the need for radiotherapy. While this trial failed to reveal any difference in overall survival (OS)^14^, a retrospective subgroup analysis from this trial reported increased OS specifically in NSCLC patients^15, 16^. In our study, we sought to investigate the involvement of RANKL in primary tumors from NSCLC using genetically engineered mouse models.

## Material and methods

### Animal models

*K-ras^LSL-G12D/WT^* (K) and *p53^FL/FL^* mice in a C57BL6/J background were purchased from The Jackson Laboratory, and were bred together to obtain *K-ras^LSL-G12D/WT^*; *p53^FL/FL^* (KP) mice. *Lkb1^FL/FL^* mice in a mixed background (FVB; 129S6) were obtained from R. DePinho (The University of Texas MD Anderson Cancer Center) through the National Cancer Institute mouse repository, were backcrossed seven times to C57BL6/J and were bred with *K-ras^LSL-G12D/WT^* mice to obtain *K-ras^LSL-G12D/WT^*; *Lkb1^FL/FL^* (KL) mice.

### Mouse treatment modalities

Mice were treated with 10 mg kg^−1^ RANK-Fc or Fc control, by subcutaneous injection twice weekly. Cisplatin was used at 3.5 mg kg^−1^ in sterile PBS by intraperitoneal injection.

### Study approval

All mouse experiments were performed with the permission of the Veterinary Authority of the Canton de Vaud, Switzerland (license number VD2391).

## Results and discussion

### RANKL expression correlates with poor overall survival and KRAS mutation in lung adenocarcinoma

To test the hypothesis that RANKL signaling regulates the development of primary lung tumors, we interrogated two independent publicly available gene expression datasets of human lung adenocarcinoma, and one of colorectal cancer (CRC). It revealed that high expression of *TNFSF11* (*RANKL*) was associated with a reduced OS compared to tumor samples with low expression in lung adenocarcinoma but not in CRC (Fig. 1A and S1A). *TNFRSF11A* (*RANK*) expression was also associated with reduced OS, but only in one of the two lung datasets, whereas it was linked with better OS for CRC patients (Fig. S1B-C), suggesting a peculiar relationship between RANK/RANKL signaling and lung cancer progression that is not shared by CRC. Specific data analysis from The Cancer Genome Atlas (TCGA, <u>https://tcga-data.nci.nih.gov/tcga/</u>) also showed that *TNFSF11* expression was significantly stronger in *KRAS* mutant compared to *KRAS* wild type tumors in lung adenocarcinoma but not in CRC (Fig. 1B). *KRAS* mutation itself in lung adenocarcinoma is not prognostic of poor OS as reported earlier^17^ (Fig. S1D). Recently, gene expression analysis from TCGA enabled to define biologically distinct *KRAS* mutant lung tumors in function of co-occurring mutations in tumor suppressors^18^. Specifically, this analysis highlighted three different subgroups, called “KC” (mutant for *KRAS* and enriched for *CDKN2A* mutations), “KP” (mutant for *KRAS* and enriched for *TP53* mutations) and “KL” (mutant for *KRAS* and enriched for *LKB1* mutations). Analogies were drawn between cluster-specific biological signatures and tumors from different mouse models of oncogenic Kras-dependent lung tumorigenesis. For example, gene set enrichment analysis (GSEA) of the “KL” cluster retrieved gene signatures obtained from murine lung tumors expressing mutant *Kras(G12D)* and deficient for *Lkb1*^18^. Intrigued by these analogies, we decided to compare *RANKL* expression between each cluster. This analysis showed that *RANKL* levels were the most elevated in the “KL” cluster, stronger (but not statistically different) than the “KC” cluster, and statistically significantly (*P* = 9.6e-07) stronger than the “KP” cluster (Fig. 1C). From each of the *KRAS* mutant and *KRAS* WT group, as well as the “KC”, “KL” and “KP” clusters from *KRAS* mutant tumors, we interrogated the *TNFSF11* most positively correlated genes (Table S1). This analysis revealed hallmark pathways of cell cycle, mammalian target of rapamycin (mTOR) signaling, epithelial-mesenchymal transition (EMT), hypoxia and glycolysis in the *KRAS* mutant samples. In contrast, *TNFSF11* correlated genes in *KRAS* WT tumor samples revealed pathways representing EMT, inflammatory and angiogenic responses and, interestingly, KRAS signaling. *KRAS* mutant sub-group analysis highlighted further differences between the clusters, with enrichments for pathways of cell cycle and EMT in “KC”, cell cycle and mTOR signaling in “KL”, and inflammatory and immune responses, KRAS signaling and EMT in “KP” (Table S1). Thus, *RANKL* expression is associated with poor OS in human lung adenocarcinoma, is stronger in *KRAS* mutant samples and the strongest when *LKB1* co-occurring mutations are present.

**Figure 1.**
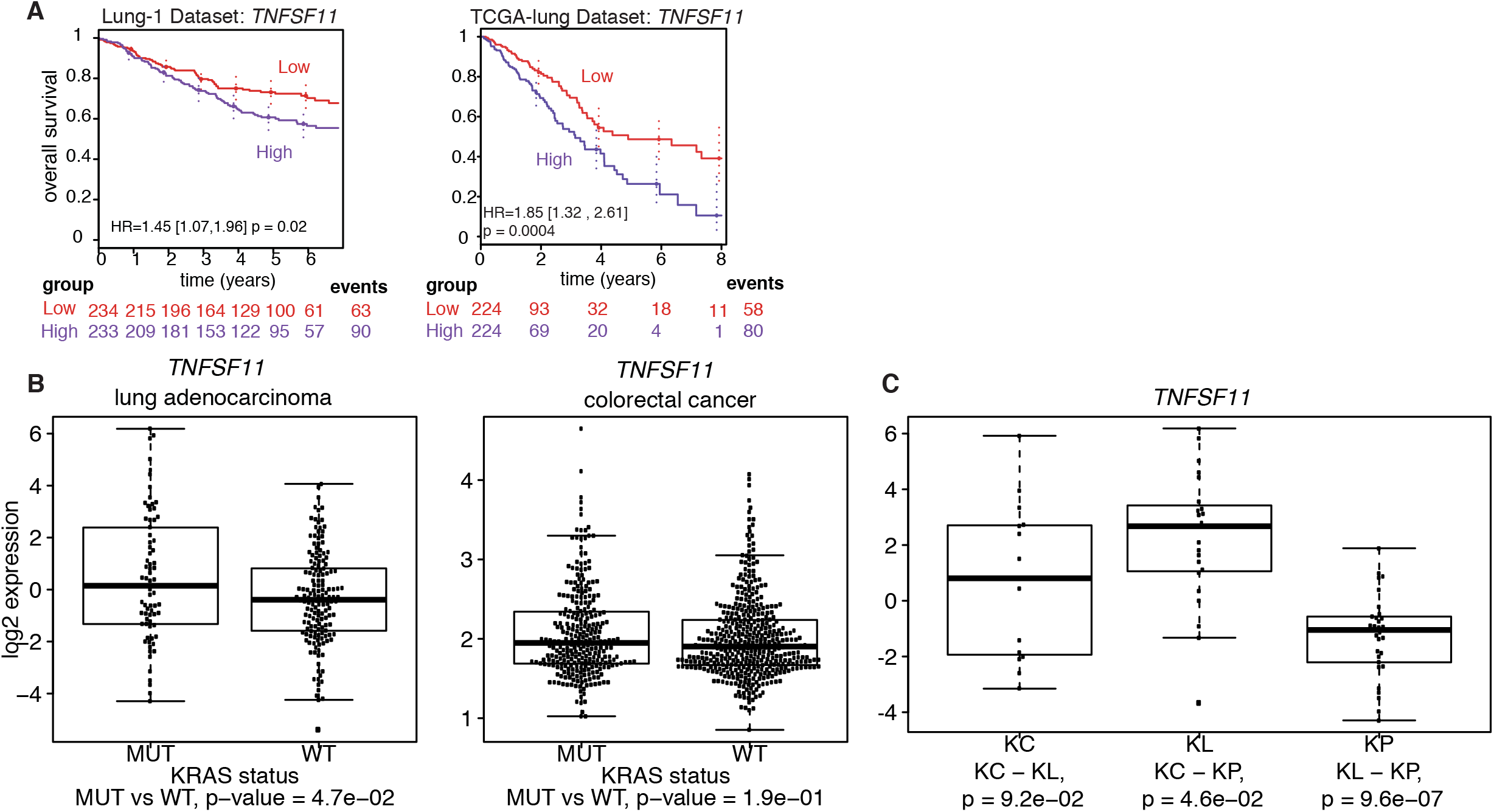
*RANKL* expression correlates with poor OS and *KRAS* mutation in human lung adenocarcinoma. (**A**) Kaplan–Meier curves for overall survival, hazard ratios (HR), confidence interval and *P*-values of pairwise differences between groups with high or low *TNFSF11* (*RANKL*) expression in (left) Lung-1 and (right) TCGA-lung adenocarcinoma datasets. Differences between curves were assessed using the log rank test (Mantel-Cox). (**B**) *TNFSF11* expression level (log2) in (left) TCGA-lung and (right) PETACC colorectal cancer (CRC) datasets split into two groups based on *KRAS* mutation status (TCGA-lung: MUT, *n*=60; WT, *n*=138; PETACC: MUT, *n*=283; WT, *n*=425). Statistical analyses were performed using Kruskal-Wallis test. (**C**) *TNFSF11* expression level (log2) in “KC”, “KL” and “KP” clusters as defined in^18^. *P*-values are indicated. Statistical analyses were performed using Kruskal-Wallis test.

### Pharmacological blockade of RANKL inhibits tumor growth in mouse models of lung adenocarcinoma

The human data prompted us to investigate the possibility that RANKL signaling contributes to primary tumor development in lung adenocarcinoma. To test this, we decided to block RANKL-RANK interaction with a recombinant protein composed of the extracellular portion of mouse RANK fused to human Fc (RANK-Fc), first in two genetically-engineered mouse models of NSCLC, *K-ras^LSL-G12D/WT^* (designated “K” thereafter) and *K-ras^LSL-G12D/WT^*; *Lkb1^FL/FL^* (“KL”). In these models, tumor initiation is triggered in adult mice upon intratracheal delivery of viruses carrying the Cre recombinase, resulting in oncogenic *Kras(G12D)* activation (K), or *Kras(G12D)* activation concomitantly with deletion of tumor suppressor *Lkb1* (KL). Loss of *Lkb1* enables tumor progression toward aggressive adenocarcinoma, with transdifferentiation to squamous cell carcinoma in some cases^19–21^. For preclinical relevance, mice were treated with RANK-Fc or control Fc only when tumors were well established and visible by micro-computed tomography (μCT). Short term (2 weeks for K, 1 week for KL mice) RANK-Fc treatment led to a significantly diminished tumor growth rate, with tumor shrinkage (i.e. relative growth below 1) detected in multiple tumors, as monitored by μCT (Fig. 2A). Next, to determine if there is rapid tumor progression following initial sensitivity to RANKL inhibition, we treated the most aggressive model twice weekly with RANK-Fc or control Fc for a total duration of three weeks, and monitored tumor growth rates longitudinally by μCT. Tumor growth remained compromised, indicating a durable response of primary lung adenocarcinoma to RANKL blockade (Fig. 2B). This response was accompanied by a reduced proliferation of tumor cells (Fig. 2C). Hence, *Kras* mutant lung tumors expressing or not *Lkb1* are sensitive to RANKL blockade.

**Figure 2.**
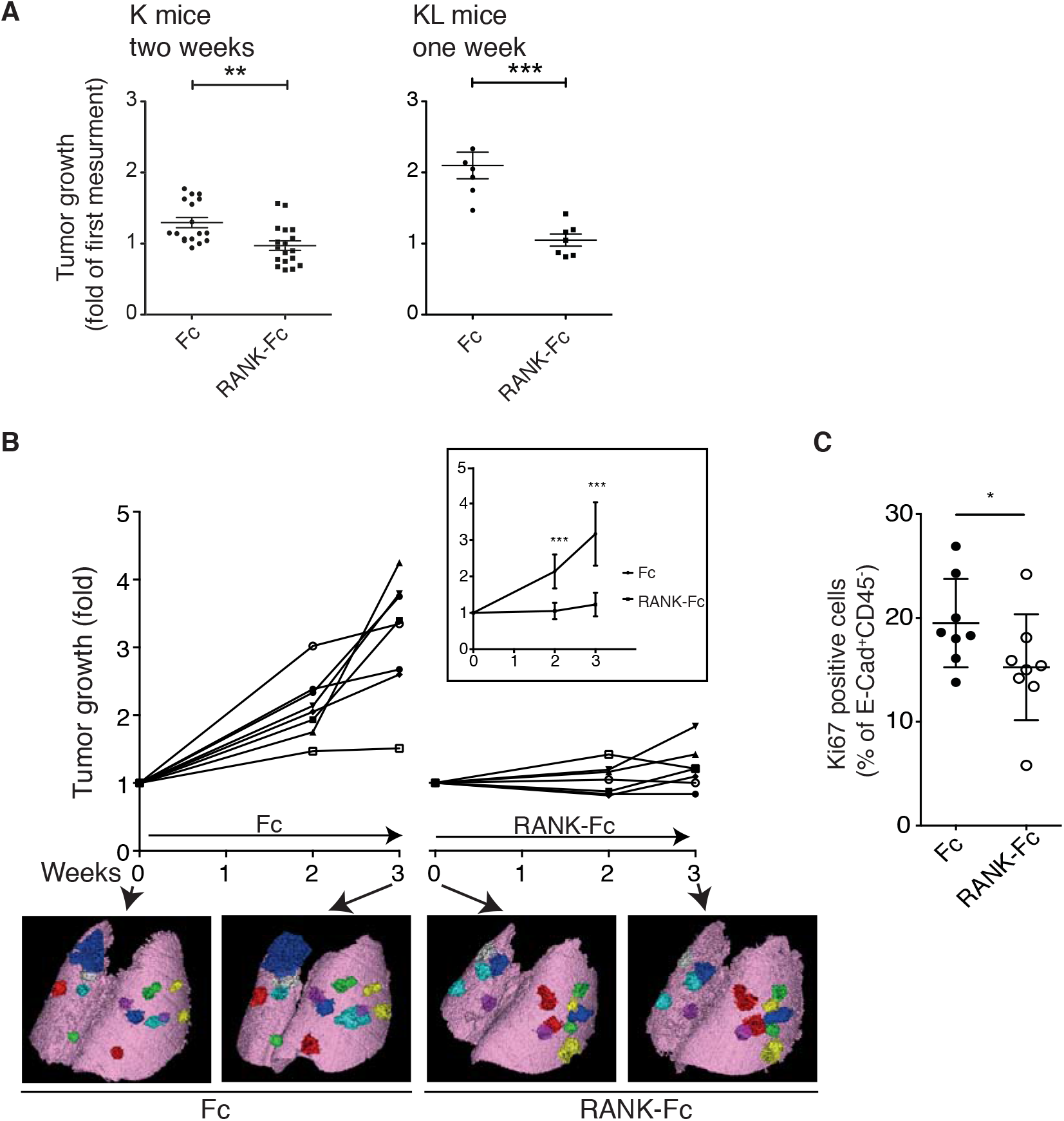
RANKL blockade inhibits the growth of lung tumors in *Kras^LSL-G12D/WT^* and *Kras^LSL-G12D/WT^*; *Lkb1^FL/FL^* mice. (**A**) Dot plots represent tumor growth, as measured by μCT, between treatment initiation and ending, in control Fc or RANK-Fc treated K (*n*=17 vs 18) and KL (*n*=7 vs 7) mouse models. Treatment started at respectively 23 and 18 weeks post-tumor initiation, and treatment duration is indicated. (**B**) Dot plot shows the percentage Ki67 positive cells among E-Caderin^+^CD45^−^ viable tumor cells from KL tumors after one week of treatment with control Fc (*n*=8) or RANK-Fc (*n*=8). (**C**) Tumor growth monitoring by μCT over a 3-week period. KL mice were treated with control Fc or RANK-Fc. 8 and 7 tumors were monitored for Fc or RANK-Fc treatment, respectively. Curves represent volume evolution of each tumor as fold of the first measurement. μCT imaging was done at weeks 0, 2 and 3 of treatment. Means with SEM are shown in the box. μCT images show lungs and the detectable tumors of two representative mice at week 0 and 3 of treatment. (**A-C**) Statistical analyses were performed using Mann-Whitney test; asterisks show *P*-values: * *P*<0.05, ** *P*<0.01, *** *P*<0.001.

### RANKL blockade elicits long term antitumor responses in K-ras^LSL-G12D/WT^; p53^FL/FL^ mice

To evaluate the antitumor effects of RANKL blockade in yet another model, we treated tumor-bearing *K-ras^LSL-G12D/WT^*; *p53^FL/FL^* (“KP”) mice^22^ with RANK-Fc. Similarly to the effect observed in K and KL mice, KP tumors were inherently sensitive to short term RANKL blockade, although no change in tumor cell proliferation could be measured contrasting with KL tumors (Fig. 3A, B). Phospho-Erk, a marker of advanced grade in *Kras^G12D^, p53*-deficient lung tumors^23, 24^, was reduced in tumors from KP mice treated with RANK-Fc (Fig. 3C). Six-week treatment follow-up indicated a long-term response to RANKL blockade in KP mice (Fig. 3D). Together, the results from Figures 2 and 3 demonstrate an inherent sensitivity and durable response to RANKL blockade in primary lung tumors in the mouse.

**Figure 3.**
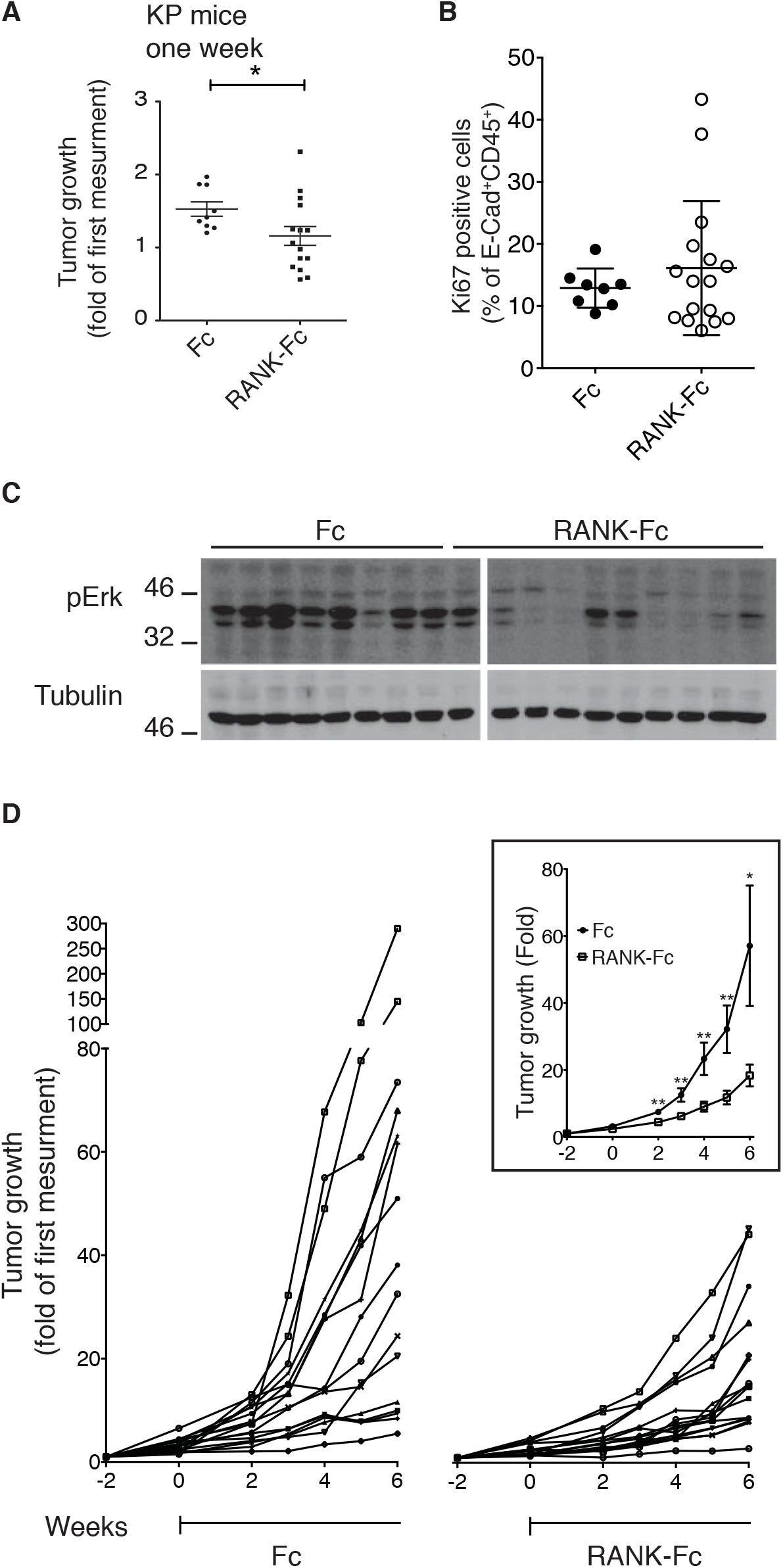
RANKL blockade elicits long-term antitumor response in *Kras^LSL-G12D/WT^*; *p53^FL/FL^* mice. (**A**) Dot plots represent tumor growth, as measured by μCT, between treatment initiation and ending, in control Fc or RANK-Fc treated KP (*n*=9 vs 15) mice. Treatment started at 13 weeks post-tumor initiation, and lasted one week. (**B**) Tumor growth monitoring by μCT over an 8-week period. KP mice were treated during the last 6 weeks with control Fc or RANK-Fc. 17 tumors were monitored for each treatment. Curves represent volume evolution of each tumor as fold of the first measurement. μCT imaging started 2 weeks before treatment initiation, and treatment started at week 0. Means with SEM are shown in the box. (**C**) Western blot was performed from KP tumor extracts after one week control Fc (*n*=8) or RANK-Fc (*n*=10) treatment, to monitor phospho (p)-Erk levels. β-tubulin was used as loading control. (**D**) Dot plot represents the percentage of Ki67^+^ cells among the E-cadherin^+^CD45^−^ cells from KP tumors after one week of treatment with control Fc (*n*=8) or RANK-Fc (*n*=16). (**A, B, D**) Statistical analyses were performed using Mann-Whitney test; asterisks show *P*-values: * *P*<0.05, ** *P*<0.01.

### RANKL-RANK expression is elevated in tumor-associated immune cells but blockade does not evoke an antitumor immune response

Next we decided to characterize better the effect of RANKL blockade, specifically in KP that represents the most common molecular subtype of the human malignancy. Staining of tumor sections by immunofluorescence enabled us to detect each of RANK and RANKL plasma membrane protein expression heterogeneously within the tumor mass, with RANK frequently co-localizing with E-cadherin (Fig. 4A). To understand which cell types express *Tnfsf11* (*Rankl*) and *Tnfrsf11a* (*Rank*) we collected lung tumors or healthy lung from KP mice, and used anti-CD45 magnetic beads to separate non-immune (CD45^−^) from immune (CD45^+^) cells. This revealed a stronger expression of *Tnfsf11* in CD45^−^ cells isolated from tumors, which comprise mostly but not exclusively tumor epithelial cells, compared to CD45^−^ cells from healthy lung, whereas *Tnfrsf11a* expression was not different (Fig. 4B). In tumors, expression of *Tnfsf11* and *Tnfrsf11a* was more elevated in immune than non-immune cells, with the strongest expression in B and T lymphocytes for *Tnfsf11*, and in myeloid cells for *Tnfrsf11a* (Fig. 4C, D). Because of this strong expression in immune cells within lung tumors, we next wanted to know if treatment causes remodeling of the tumor immune microenvironment. RANK-Fc led to a significant increase in *Tnfsf11* expression in non-immune cells, and a reduction in immune cells, whereas there was no perturbation in *Tnfrsf11a* expression (Fig. 4E). After two weeks of treatment, no change was observed in the proportion of B cells, whereas that of T lymphocytes diminished, which was not specific for any of the CD4^+^, Treg or CD8^+^ subsets (Fig. S2A-C). RANKL blockade did not result in any significant change in the expression of *Tnf, Il10* or *Ifng* mRNAs in CD45^−^ or CD45^+^ cells, and did not alter cytoplasmic expression of IFNγ or TNF proteins in CD4^+^ or CD8^+^ T cells (Fig. S2D and E). RANK was originally characterized in DCs and suggested to participate in DC-dependent T cell expansion, a hypothesis that was not supported by analysis of RANK deficient mice^3, 4^ Our results demonstrate that, despite elevated RANK and RANKL expression in immune cells within the tumor microenvironment, RANKL blockade only marginally remodels the immune compartment, which does not support a major role for immune cells in the antitumor response.

**Figure 4.**
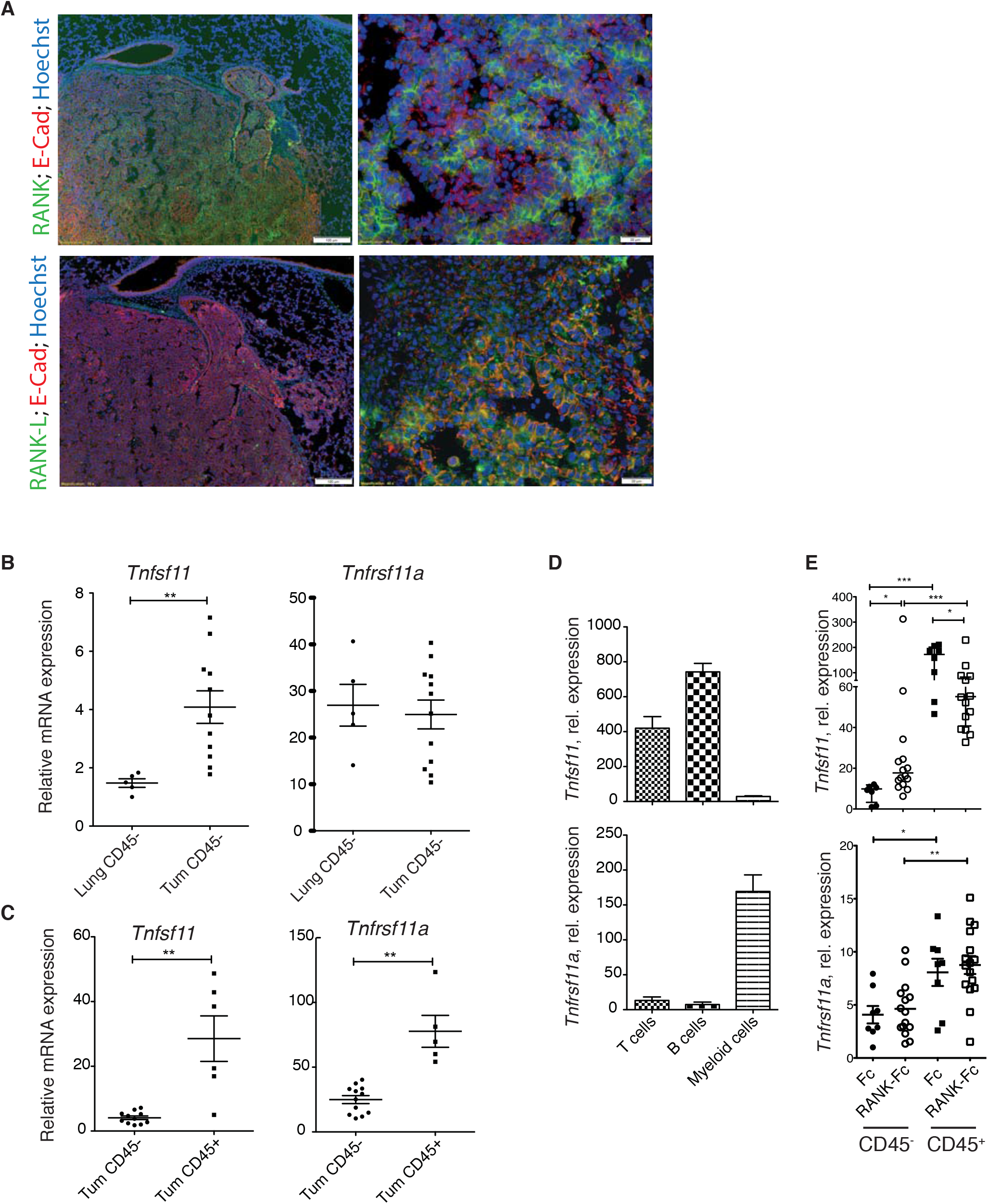
Tumor and immune cells differentially express RANKL and RANK. (**A**) Representative immunofluorescence staining of a KP tumor showing nuclei (Hoechst, blue), E-cadherin (red) and (upper panels) RANK or (lower panels) RANKL (green). Scale bars are 100 μm (left) or 20 μm (right). (**B**) *Tnfsf11* (*Rankl*) and *Tnfrsf11a* (*Rank*) relative mRNA expression from non-immune (CD45^−^) cells of normal lung (n=5) or KP tumors (Tum, *n*=11). (**C**) *Tnfsf11* and *Tnfrsf11a* relative mRNA expression of sorted CD45^−^ (*n*=11) or CD45^+^ (*n*=6) cells from KP tumors. (**D**) *Tnfsf11* and *Tnfrsf11a* relative mRNA abundance in sorted CD3^+^ T cells, B220^+^ B cells and CD3^−^B220^−^CD45^+^ myeloid cells from KP tumors. Histograms represent the average of multiple experiments and error bars denote SEM. *n*=4, 4 and 7 experiments respectively for T, B and myeloid cell populations. (**E**) *Tnfsf11* and *Tnfrsf11a* relative mRNA expression in CD45^−^ and CD45^+^ cells of tumors from control Fc or RANK-Fc treated KP mice. Treatment started at 23 weeks post-tumor initiation, and duration was 2 weeks. (**B, C** and **E**) Statistical analyses were performed using Mann-Whitney test. * *P*<0.05, ** *P*<0.01, *** *P*<0.001.

### RANKL blockade increases the efficacy of cisplatin chemotherapy

To test the effect of combination treatment compared to monotherapy, we decided to treat tumor-bearing KP mice with RANK-Fc for one week followed by one week of standard chemotherapy cisplatin with maintenance of RANK-Fc, and used μCT to compare tumor growth rates to that of tumors from mice treated with cisplatin alone. Short-term cisplatin caused a diminished tumor growth rate, as reported earlier^25^, which was not significantly different from RANK-Fc therapy (Fig. 5A). Combined RANK-Fc + cisplatin led to a more profound effect on tumor growth (Fig. 5B). When compared to cisplatin alone, combined therapy led to a significant decrease in the expression of the anti-apoptotic genes *Bcl2l1* and *Xiap* in the tumor cell compartment, as well as *Xiap* in immune cells (Fig. 5C). Hence, our results highlight the efficacy of RANKL blockade in preclinical mouse models of lung adenocarcinoma. They suggest different mechanisms of action in different tumor molecular subtypes, and that combination treatments will be more efficacious than chemotherapy alone in patients (Fig. 5D).

**Figure 5.**
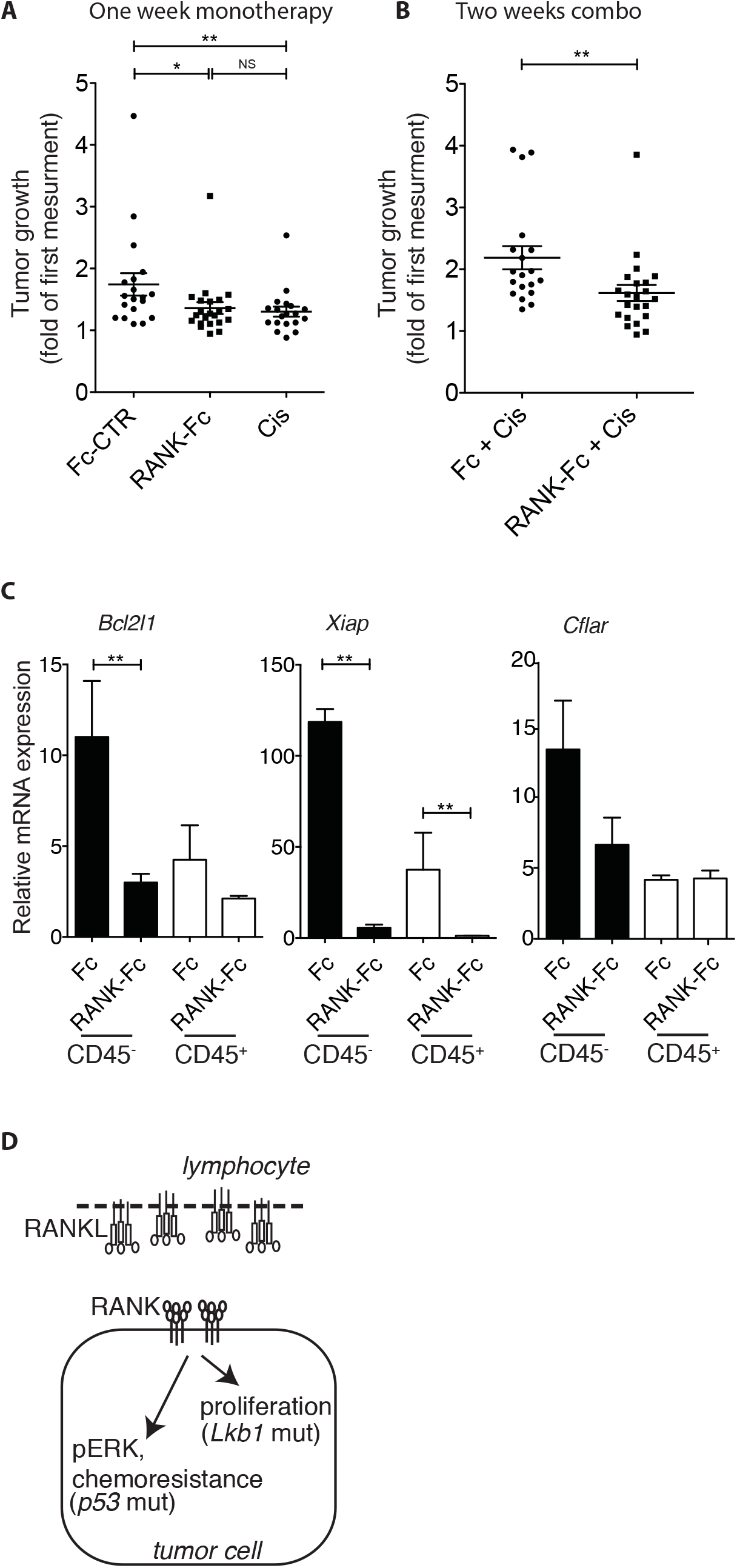
RANKL blockade increases chemotherapy efficacy. (**A**) Dot plot shows tumor growth, determined by μCT imaging, in KP mice treated or not during one week with Fc control (Fc-CTR) (*n*=19), cisplatin (Cis) (*n*=19) or RANK-Fc (*n*=22). (**B**) Dot plot shows tumor growth, determined by μCT imaging, in KP mice treated by a combination of control Fc (*n*=19) or RANK-Fc (*n*=22) plus cisplatin. Mice received Fc or RANK-Fc injections during two weeks plus one injection of cisplatin during the last week. (**C**) Relative mRNA abundance of the indicated genes in CD45^+^ or CD45^−^ sorted cells of tumors obtained from KP mice treated in (**B**). Error bars show SEM. Statistical analyses comparing CD45^+^ versus CD45^−^ fractions were not performed. (**A-C**) Statistical analyses were performed using Mann-Whitney test; Asterisks show statistical *P*-values: * *P*<0.05, ** *P*<0.01, NS, not significant.

We have used mouse models where tumor initiation depends on oncogenic *Kras* activation, because (1) *KRAS* is the most commonly mutated proto-oncogene in human NSCLC, (2) these models faithfully recapitulate many aspects of the development of the human disease, and (3) these models are refractory to many treatments and are thus relevant to assess new therapeutic opportunities (here RANKL blockade). Our preclinical results, combined with our bioinformatics analyses, depict a strong connection between mutant *KRAS* and RANKL signaling in lung adenocarcinoma, which probably does not take place in CRC. Hence our observations are generating the hypothesis that RANKL blockade will be efficient in *KRAS* mutant lung tumors. An important future step will be to verify this hypothesis directly in tumor samples from patients.

### Case report of a patient with stage IV NSCLC treated with denosumab monotherapy

A female patient, former smoker at 30 pack-year, presented with a middle lobe lung adenocarcinoma, *EGFR* wild-type, without *ALK* rearrangement, cT4 cN0 cM1b (lower right lobe and bone metastases in the thoracolumbar spine especially L3, pelvis, femoral collar) stage IV. She was diagnosed on 15.03.2013 (transbronchial biopsy lower right lobe). The patient was hesitating regarding chemotherapy. While she was presenting several painful bone metastases, an initial treatment of denosumab monotherapy was introduced as the first step in her oncology management, in parallel to supportive care including lumbar spine palliative radiotherapy in March. She decided to start chemotherapy on the 21^st^ of May, 2 months after diagnosis. A new CT scan was performed before starting chemotherapy, 4 weeks after a unique cycle of denosumab only, showing a primary lung tumor partial response (Fig. 6A and B). Thereafter she received systemic chemotherapy (10 cycles, 4 platinum-pemetrexed, 6 pemetrexed, with partial response and thereafter with stable disease). She died 7 months after diagnosis.

**Figure 6.**
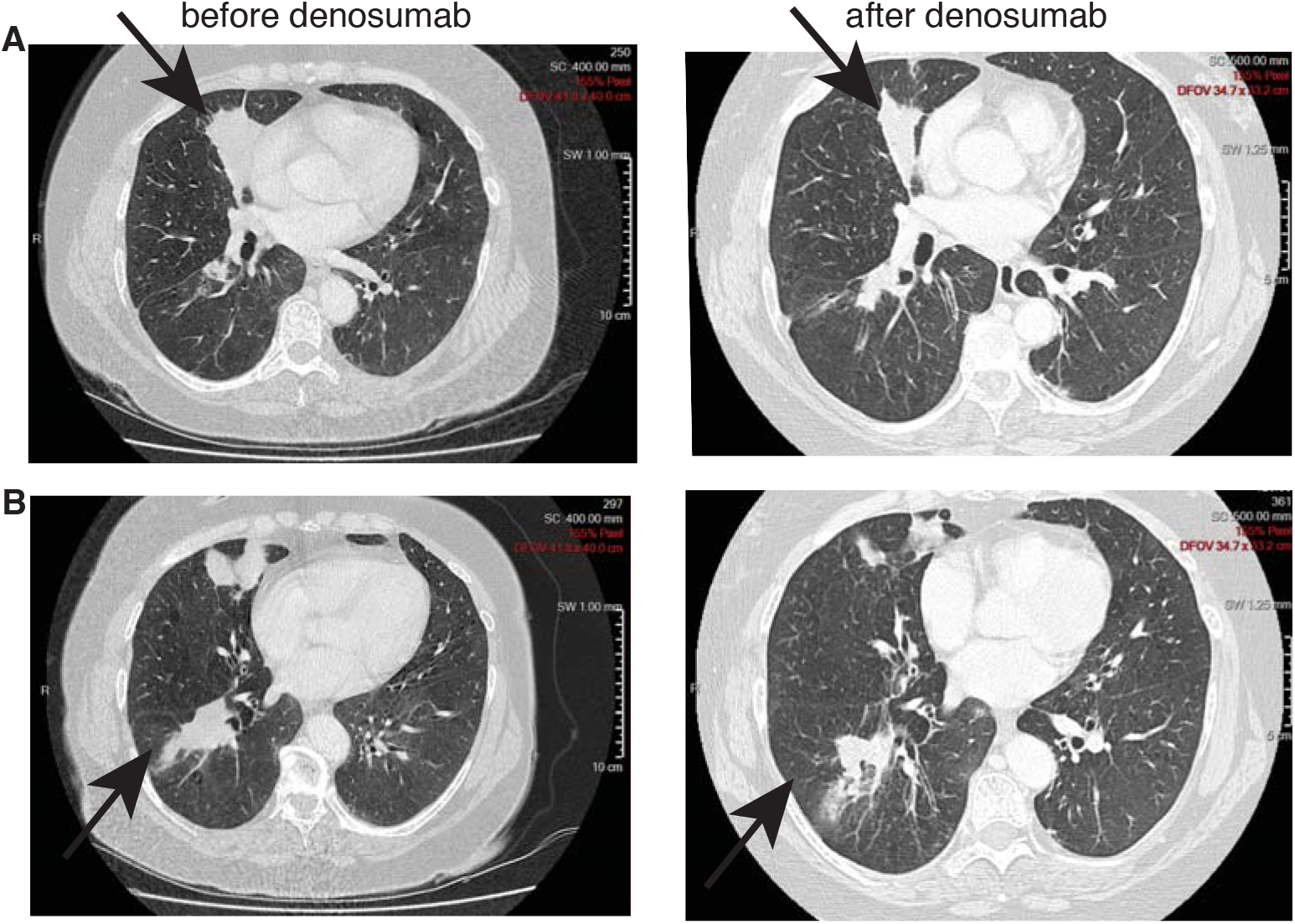
Clinical efficacy of denosumab monotherapy in a NSCLC patient. (**A, B**) CT scans of the thorax of a chemotherapy naïve stage IV NSCLC patient showing (**A**) middle lobe adenocarcinoma and (**B**) a lower lobe secondary lesion before (left) and after (right) a single dose of denosumab, demonstrating a significant partial response 4 weeks after a single infusion.

This clinical case illustrates the potential efficacy, at least in select cases, of RANKL blockade therapy even in advanced disease as monotherapy. Furthermore, the very large unplanned retrospective analysis of denosumab versus zoledronic acid in bone metastatic lung cancer^16^ as well as our gene expression dataset analyses of human lung adenocarcinoma both support such hypothesis of an improved OS. The activity of denosumab as an anti-neoplastic drug for the treatment of advanced NSCLC in combination with frontline platinum-based chemotherapy is currently being evaluated in a large randomized phase 3 EORTC/ETOP trial SPLENDOUR, with a primary endpoint of OS, and a parallel retrospective analysis of RANK signaling pathway activation, including RANK and RANKL expression at baseline for all patients.

## Author Contributions

Conception and design of the project: JF, SP and EM; *In vivo* experiments: JF and CC; bioinformatics analyses: NZ; Data analysis: JF, CC, SP and EM; Description of case report: SP; Project supervision: EM; Manuscript writing: JF and EM. All authors discussed the results.

## Acknowledgments

We thank the EPFL SV Histology Core Facility for histological sectioning, and the EPFL SV Flow Cytometry Core Facility. We thank E. Kadioglu who helped in mouse cohort generation, H. Ramay for the initial help with bioinformatics analyses, H. Golay and J. Vazquez for technical assistance, and M. Pittet for critical reading of the manuscript.

